# Pharmacogenetic manipulation of Locus coeruleus activity in monkeys demonstrates its role in cognitive effort

**DOI:** 10.64898/2026.07.23.740243

**Authors:** Pauline Perez, Sebastien Bouret

## Abstract

The noradrenergic nucleus locus coeruleus (LC) is involved in numerous cognitive functions. Its activation often enhances sensory and motor performance and its activity correlates with arousal, which altogether suggest a general role in the mobilization of resources for cognition and action. We recently showed its strong and specific implication in physical effort and indirect evidence suggest that it might also be involved in cognitive effort. To address this question directly, we used a pharmacogenetic approach in rhesus macaques to selectively and reversibly inhibit LC neurons in a cognitively challenging task. Two monkeys were injected with viral vectors expressing inhibitory DREADDs (hM4Di) specifically in noradrenergic LC neurons, allowing reversible suppression of LC activity via systemic administration of deschloroclozapine (DCZ, 0.1 mg/kg). A third monkey served as a control and only received DCZ injections. Monkeys performed a simple hole-board task in which they searched for food rewards (raisins) hidden in a 5×5 grid of wells. In the transparent condition, rewards were visible, requiring minimal cognitive effort. In the opaque condition, rewards were hidden, such that monkey had to rely upon working memory to avoid revisiting empty wells. Thus, performance in opaque condition required more cognitive effort. Behavioral analysis showed that LC inhibition had no effect on performance in the transparent condition. However, in the opaque condition, it significantly impaired performance by increasing errors (revisits), without affecting the total number of rewards obtained, response times, or overall motivation. Monkeys compensated the decrease in success rate by performing more trials, indicating reduced efficiency rather than disengagement. These findings demonstrate that the LC plays a critical causal role in mobilizing cognitive resources for a demanding task. This is in line with the idea that the noradrenergic system contributes broadly to cognitive control and effort, extending previous findings on its involvement in physical effort. Overall, the study provides strong evidence linking LC activity to cognitive effort regulation, thereby complementing non-invasive studies in humans.

## Introduction

The noradrenergic nucleus locus coeruleus (LC) has long been associated with cognition. After early studies identified its strong relation with arousal and vigilance, several authors suggested that noradrenaline (NA) could also play a critical role in executive functions (Berridge and Waterhouse 2003, Poe, Foote et al. 2020). For example, NA has a positive influence on working memory (Arnsten 2000, Rossetti and Carboni 2005, Hernaus, Casales Santa et al. 2017). Manipulations of the NA system were also shown to affect attention (Witte and Marrocco 1997, Riekkinen, Kejonen et al. 1998, Reynaud, Froesel et al. 2019), as well as inhibitory control (Chamberlain, Hampshire et al. 2009, Rae, Nombela et al. 2016).

The activity of LC neurons strongly correlates with autonomic arousal, (Abercrombie and Jacobs 1987, Jacobs, Abercrombie et al. 1991, Aston-Jones, Rajkowski et al. 1996). Based on these findings, several authors used pupil diameter as a proxy for LC activity to study its role in cognitive functions in humans. This line of work provided a clear insight into the cognitive processes associated with transient increases in arousal, and potentially involving the LC (Einhäuser, Stout et al. 2008, Nassar, Rumsey et al. 2012, de Gee, Tsetsos et al. 2020, Filipowicz, Glaze et al. 2020). Several imaging studies confirmed the relation between the activation of the LC and pupil dilation, and proposed an interpretation in terms of mobilization of cognitive resources (Alnaes, Sneve et al. 2014, Murphy, O’Connell et al. 2014, Tomassini, Hezemans et al. 2022, Negelspach, Alkozei et al. 2025). Indeed, earlier studies had identified the strong relation between autonomic arousal (as measured using pupilometry) and the mobilization of cognitive effort (Kahneman and Beatty 1966, Pribram and McGuinness 1975). More recently, pupil dilation was associated with cognitive control, broadly defined as a mobilization of cognitive resources to overcome mental challenges, which is analogous to the notion of mental effort (Botvinick, Braver et al. 2001, Naccache, Dehaene et al. 2005, Howells, Stein et al. 2010, Alnaes, Sneve et al. 2014, Lisi, Bonato et al. 2015, Zenon, Sidibe et al. 2015, Shenhav, Musslick et al. 2017, van der Wel and van Steenbergen 2018, Unsworth, Miller et al. 2022).

This relation between pupil and cognitive effort could be related to our previous studies showing the strong relation between the LC/NA system and effort in rhesus monkeys. Indeed, we first showed that both the timing and the magnitude of LC activation were related to the production of physical effort, as well as the associated pupil dilation (Varazzani, San-Galli et al. 2015). Pharmacological manipulations demonstrated that NA was causally involved in physical effort, in that decreasing NA levels with clonidine induced both an increase in effort costs (for decision-making) and a decrease in force production (Borderies, Bornert et al. 2020). Finally, we also showed that transient LC activations were observed when monkeys were producing a mental effort (Bornert and Bouret 2021). Thus, it is tempting to propose that the LC/NA systems plays a general role in effort, both cognitive and physical. Indeed, an interpretation in terms of cognitive effort could also capture the known influence of NA manipulations on performance in tasks requiring working memory, attention or inhibitory control (Chamberlain, Müller et al. 2006, Robbins and Arnsten 2009, Reynaud, Froesel et al. 2019). But even if that hypothesis seems plausible given the strong relation between effort, LC activity and autonomic arousal (as index using pupil dilation), this relation is far from being specific: numerous other brain regions are associated with cognition and autonomic arousal (Saper 2002, Barbas and Saha 2003, Critchley 2009, Joshi, Li et al. 2016, Reimer, McGinley et al. 2016). Thus, even if the implication of the LC in cognitive effort seems likely, its relative contribution still needs to be quantified, and directly evaluated using causal manipulations. Given the complexity of NA pharmacology, with multiple receptors subtypes located not only in distinct parts of the brain but also in the periphery, interpreting pharmacological manipulations has always been challenging (Berridge and Waterhouse 2003, Robbins and Arnsten 2009, Poe, Foote et al. 2020). Interestingly, recent pharmacogenetic studies in rodents provided critical insight on LC functions, by enabling a targeted and reversible manipulation of LC activity (Hirschberg, Li et al. 2017, Zerbi, Floriou-Servou et al. 2019, Bacon, Pickering et al. 2020, Cerpa, Piccin et al. 2023, Xia, Maheu et al. 2026). Thus, our goal here was to take advantage of that approach to study the role of LC in cognitive effort in monkeys.

In this study, we explored the causal role of the LC in cognitive effort by evaluating the influence of targeted and reversible inhibition of LC neurons in rhesus macaques performing a simple working memory task. We used a pharmacogenetic approach to induce a specific and reversible inactivation of LC neurons using Designer Receptor Exclusively Activated by Designer Drugs (DREADD) (Perez, Chavret-Reculon et al. 2022). In our conditions, the activation of inhibitory DREADDs with DCZ (0.1 mg/kg) induced a reliable 15% decrease in the firing of LC neurons, which did not alter vigilance (Perez, Chavret-Reculon et al. 2022). We used a task previously shown to engage working memory and rely upon the integrity of the dorsal prefrontal cortex (Collin, Cowey et al. 1982, Passingham 1985). This task involves a well-board consisting of 25 wells (5 x 5), each filled with a food reward (typically, a raisin). Monkeys only need to displace a revolving door to retrieve the reward, which they do spontaneously. In control sessions, the doors are transparent and monkeys can visually keep track of already visited wells and select those still containing a reward. In Opaque condition, doors are opaque such that monkeys need to use working memory to keep track of recently visited wells, to minimise revisits and collect rewards efficiently. Once they have learned to displace the doors to collect the reward in the corresponding well, they spontaneously adjust their behavior to minimise revisits, such that the engagement of working memory in the opaque condition does not require any training. Thus, comparing the influence of LC inactivation in opaque vs transparent conditions provides a simple, yet reliable way to evaluate the implication of LC in working memory, and the corresponding engagement of cognitive effort.

## Materials and methods

### 1. Animals

All procedures complied with the guidelines of the European Community for the care and use of laboratory animals, applying the principle of 3Rs and in accordance with the welfare of the animals. They were conducted in agreement with European Community regulations (European Union Directive 2010/63/UE) and approved by the local Darwin Ethical and National committees (authorization #14611-2017122009291145 v4, 18/01/2019). These experiments involved 3 rhesus macaques (*Macaca mulatta*): R—male, 8 years old (9,5 kg); JF—male, 8 years old (19 kg); and JS—female, 8 years old (10 kg). They were hosted at the ICM primate facility, with free access to food and water for the entire duration of the experiment. Lights followed a 12h cycle with lights progressively turned on at 8.00 AM. All testing was performed in the housing cages. All 3 monkeys had already been involved in behavioral experiments before the present study.

### 2. Surgery

Both monkey JF and JS had already been injected with constructs allowing the expression of inhibitory DREADDs (hM4Di) in the locus coeruleus (LC) (Perez, Chavret-Reculon et al. 2022). The details regarding the production of the viral construct, as well as the surgical procedures, are described in detail in Perez *et al*. (2021). Briefly, we used lentiviral expression vectors carrying the hM4Di gene and a TH-promoter sequence, such that the expression was restricted to the noradrenergic neurons of the locus coeruleus. The location of the LC was first identified using Magnetic Resonance Imaging (MRI) scanning, and a first surgery was performed to place a head fixation post and recording chambers on top of the skull, above the LC on each hemisphere. A second MRI scan was performed to identify the locations at which the injector needed to be positioned to reliably target the LC, using stereotaxic grids. The intracerebral injections were completed during a second surgery during which the injector was introduced at identified positions (using the stereotaxic grids) and descended through the brain, to reach a target point located 1mm above the LC. We performed 4 injections per animal, 2 per hemisphere, separated by approximatively 2 mm in the Antero-Posterior axis. As shown in Perez et al (2021), This procedure ensures an efficient expression of hM4Di DREADDs in the LC, since DREADD activation by systemic DCZ injection induces a dose-dependent decrease in LC activity, and a decrease in vigilance at high doses, in line with known effects of pharmacological LC inhibition using clonidine (Bouret & Richmond, 2009; Cedarbaum & Aghajanian, 1976; Starke & Montel, 1973).

### 3. Behavioral task

The apparatus, similar to Collin *et al*. (1982), consisted of a grid of 5x5 cylindrical wells, each containing a food reward (a raisin). As shown on figure 1, each well was covered by a swinging door that was either transparent or opaque. The doors were made so that the monkey could reveal the content of each well by laterally pushing them, but they went back in position when released. Animals were isolated for every session of the training and testing phases.

**Figure 1:**
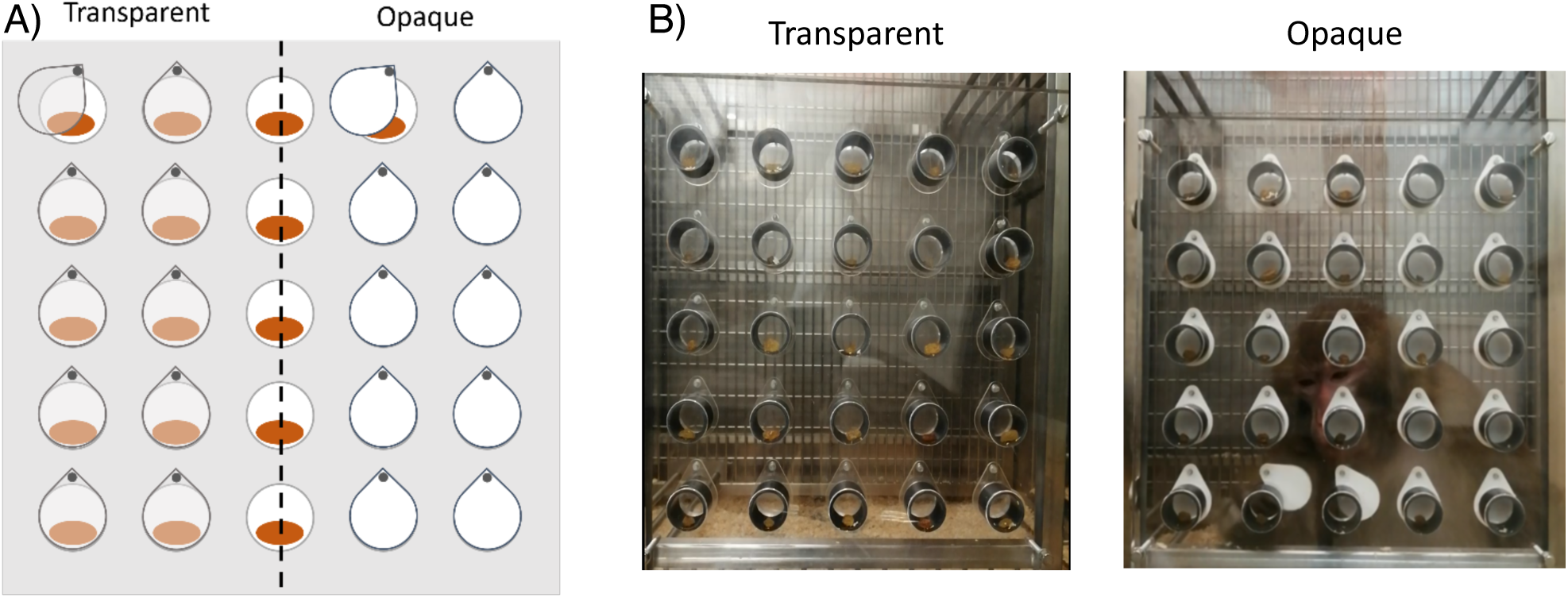
Representation of the apparatus fully baited. A) Schematic representation of the apparatus from the perspective of the monkey, in transparent (left) and opaque (right) condition. The wells on the midline (broken line) are represented with doors opened. B) Photographs of the apparatus from the perspective of the experimenter.

The training phase allowed the animals to familiarize themselves with the apparatus and to learn how to open the doors. The apparatus was placed fully baited on the cage. Different types of food rewards were tested to find the most incentive one. During this phase, the grid contained a combination of open wells, transparent doors, and opaque doors. Monkeys had free access to the apparatus. They were required to retrieve at least 23 rewards in three consecutive sessions to move on to the testing phase. In total, each animal completed six training sessions before entering the test phase.

During the testing phase, the apparatus was placed in the home cages for a maximum of 5 min, during which the monkey could freely explore the grid. If the monkey stopped interacting with the apparatus for more than an minute, the session ended and the apparatus was removed. Monkeys were tested only once a day with either only transparent or only opaque doors. In transparent condition, rewards were visible through the doors so only visual control was necessary to complete the task. In opaque condition, the rewards were no longer visible therefore, a cognitive effort was required to avoid resampling a previously visited well. Thirty minutes before the test, monkeys were injected intramuscularly with either 0.1mg/kg of DCZ diluted in NaCl or the equivalent in volume of saline solution (NaCl). Because of other experimental constraints, DCZ and saline injections could not be alternated during the first two weeks of the protocol (Figure 2). Consequently, all testing sessions during this initial period were conducted under saline, followed by two weeks of consecutive DCZ sessions. To account for a potential buildup effect of the DCZ injections, we alternated DCZ and saline injections during the last two weeks of the experiment. Transparent and opaque sessions were also not alternated during the first four weeks of the protocol, but they were semi-randomly alternated during the last two weeks of the experiment. This design resulted in a total of 14 sessions of each treatment, 13 transparent sessions and 15 opaque sessions (Table 1).

**Figure 2:**
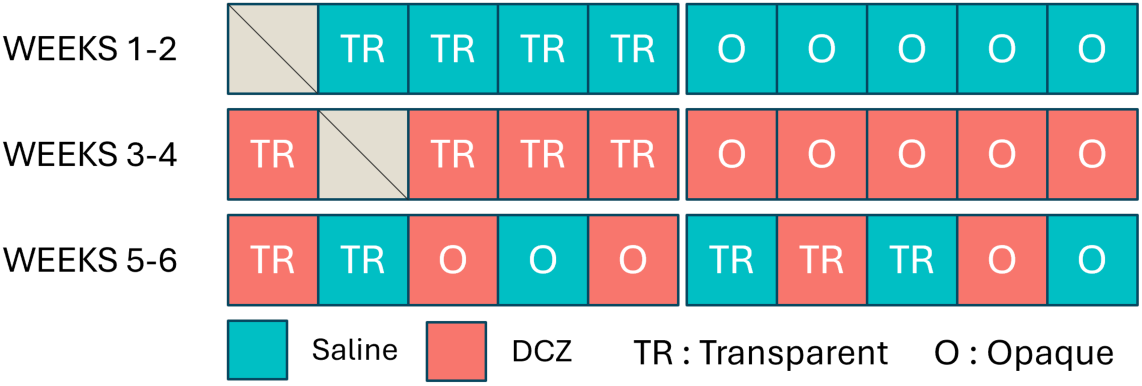
Timeline of the experiment. The first four weeks of the experiment were not randomized because of experimental constraints, while the last two were pseudo-randomized to match the number of opaque, transparent, saline and DCZ sessions.

**Table 1:** Total number of sessions in each condition, all monkeys included.

|  | Saline | DCZ | TOTAL |
| --- | --- | --- | --- |
| Transparent | 7 | 6 | **13** |
| Opaque | 7 | 8 | **15** |
| **TOTAL** | **14** | **14** |  |

### 4. Scoring and analysis

All sessions were video recorded, and the videos were scored using BORIS software (*Friard, O. and Gamba, M., BORIS: a free, versatile open-source event-logging software for video/audio coding and live observations. (2016) Methods Ecol Evol, 7: 1325–1330*.). We measured the position and timing of the choices, and whether the choice resulted in a success (a reward was retrieved), a revisit (return to a previously visited well) or a fail (a door was opened without retrieval of the reward).

All statistical tests were run with R (R Core Team, 2025. _R: A Language and Environment for Statistical Computing_. R Foundation for Statistical Computing, Vienna, Austria). Data was analyzed intra-individually and conditions were compared using t-tests and ANOVAs.

## Results

### 1) Behavior in the task under saline

#### Opaque vs Transparent

We first examined how monkeys performed in transparent (TR) condition compared to opaque (O) condition under saline treatment. The number of rewards retrieved was similar between the two conditions (T-tests, Jf: 25+/- 0, vs 24+/- 0.5 in TR vs O conditions, respectively, t(5)=1.94, p=0.11; Js: TR: 24 +/- 0.5; O: 23.1 +/- 0.7; t(11.44)=1, p=0.34; R: T: 25 +/-0, O: 24+/- 0.7; t(6)=1.45, p=0.20). The success rate (computed as the number of rewards retrieved divided by the total number of trials), however, was significantly lower in opaque condition (Figure 3A, Jf: t(10.9)=8.9, p=2.4e-06; Js: t(9.0)=2.9, p=0.02; R: t(9.9)=5.5, p=0.0003). Because the number of successes did not differ significantly between TR and O conditions, this effect was mainly driven by the total number of trials performed. Indeed, monkeys required more trials to reach the last reward of the session in O condition (Figure 3B, T-test: Jf: t(5.6)=2.6, p=0.04; Js: t(9.67)=-2.27, p=0.047; R: t(7.72)=-4.49, p=0.0022) and also performed more trials after obtaining the last reward (Figure 3C, Jf : t(5.4)=4.9, p = 0.003; Jf: W = 35, p = 0.20. R: t(7.22) = 2.26, p = 0.057). Note that similar results were obtained using non-parametric Wilcoxon signed-rank tests (supplementary material: ‘Supplementary analysis 1’).

**Figure 3:**
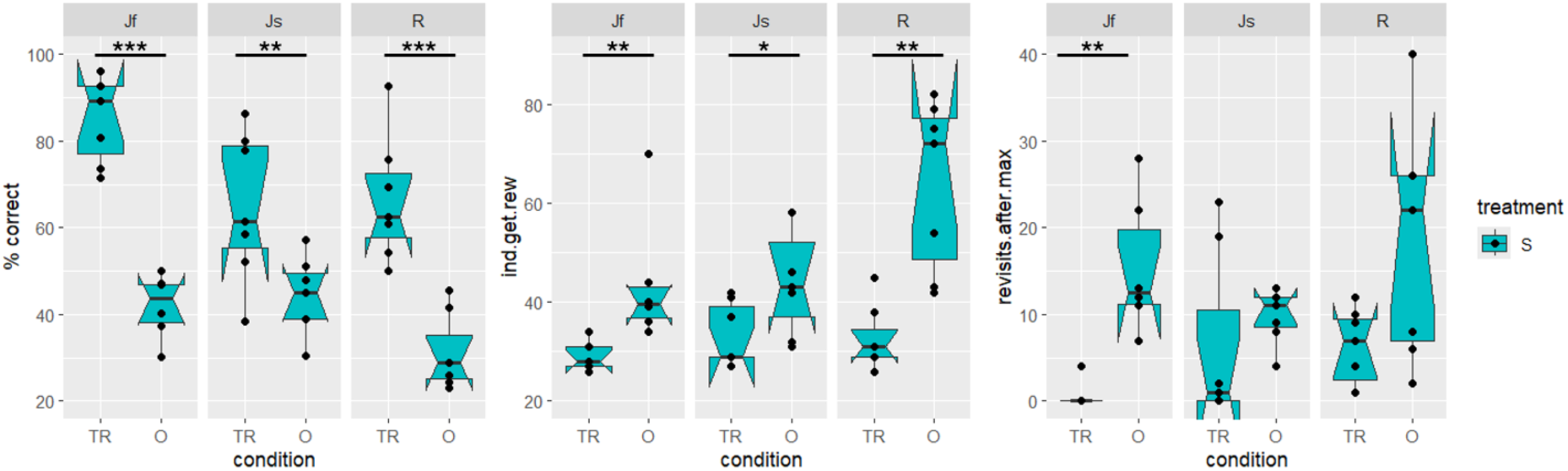
**A**. Success rate in Transparent and Opaque saline sessions. **B**. Trial number at which the last reward is retrieved in Transparent and Opaque saline sessions. **C**. Number of trials after retrieving the last reward in Transparent and Opaque saline sessions.Points represent data from individual sessions.

#### Dynamics of the performance

To capture the dynamics of each animal’s average performance across successive trials of multiple sessions, we fitted each animals’ trial-wise performance through a sigmoid function. We compared these values to the performance of two reference agents: an optimal agent and a random agent. We reasoned that an optimal agent should retrieve all 25 rewards without revisiting any position and then stop searching. The performance of a random agent is harder to evaluate intuitively, because the probability of finding a reward « by chance » decreases as animals collect more rewards. Thus, we used computer simulations to generate the behavior of a random agent in a single session. That agent selected randomly one of the 25 positions on each trial, irrespectively of past choices. Since random agents have no reason to stop generating actions, the number of trials was set arbitrarily, by matching it with the number of trials performed by the monkey in the corresponding ‘real data’ session. For each monkey, we ran 10 simulations for each of the real sessions (here, in opaque-saline condition). As shown on figure 4, monkeys’ performance was intermediate between that of the optimal agent and that of an average random agent. For all 3 monkeys, performance in O sessions was greater than chance level, as generated by random agent simulations, especially at the beginning of the sessions. Also, even if performance was much higher in TR condition than in O condition, it was still not optimal, in that monkeys still visited empty wells even when they had direct visual information.

**Figure 4.**
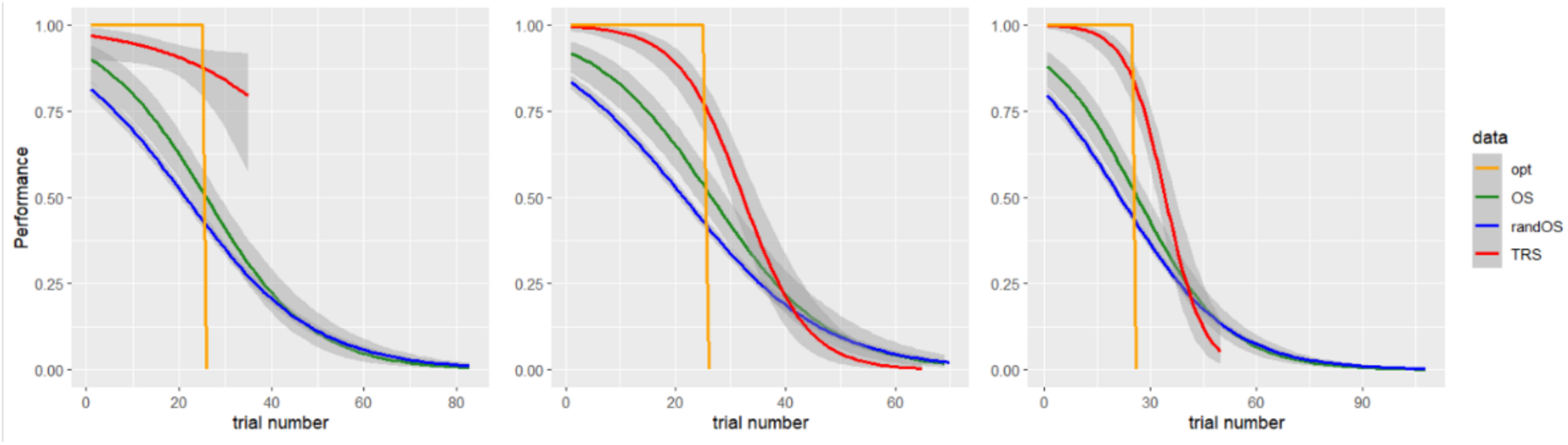
: Performance trial by trial of monkey Jf, Js and R (in order) fitted through a sigmoid function in Transparent saline condition (red), in Opaque saline condition (green), for the simulated optimal agent (orange) and the simulated random agent (blue).

At the session level, monkeys’ performance in opaque condition did not differ from the random agent simulations (T-tests, Jf: t(5.8) = 0.77, p = 0.47; Js: t(6.6) = 0.6, p = 0.6; R: t(7.2) = 0.04, p = 0.97). This is expected since, as shown on figure 4, the behavior of real animals mostly differs from that of random agents at the beginning of the sessions, when rewards are still available. Beyond 25 trials, after most rewards have been retrieved, the performance of monkeys and random agents converges to very low values. When all rewards have been collected, monkeys can only make erroneous responses and performance decreases monotonically with the number of trials. Since the number of trials in simulated trials was matched to that of real data, performance converges to similar low values, especially in opaque condition where monkeys performed more trials (see Figure 3C). Thus, we focused on the beginning of the sessions, and selected trials up to the obtention of the 20th reward (the smallest number of rewards retrieved in any session across the dataset, such that all sessions could be included). We used ANOVAs to examine the influence of two factors on performance: data type (real data vs random agents) and condition (O vs TR). The interaction between these two factors was significant for every monkey (ANOVAs, Jf: F(1,139) = 27.98, p = 3×10⁻⁷; Js: F(1,150) = 66.34, p = 1.36×10⁻¹³; R: F(1,150) = 81.3, p = 8.49×10⁻¹⁶). Indeed, in all 3 monkeys, the performance in TR was significantly better than in O, and monkeys were always performing significantly better than the simulations (PostHoc Tukey tests, p<0.05) (Figure 5A). As expected, performance was undistinguishable between O and TR conditions for random simulations, when correcting for the number of trials. Thus, even at the beginning of the sessions, monkeys performed worse in opaque than in transparent condition, but better than random agents, in line with the idea that the task was cognitively more challenging. Note that similar results were obtained using non-parametric Wilcoxon signed-rank tests (supplementary material: ‘Supplementary analysis 2’). In short, this dynamic analysis reveals that when performing the task in opaque condition, *i.e.* in absence of visual information regarding reward availability, performance is lower compared to TR condition for two reasons: 1) the animals had more difficulty to collect the rewards from the beginning of the sessions and 2) the animals persisted in performing the task longer after all rewards had been collected. Critically, these two measures did not seem to be related: in all 3 monkeys, the correlation (across sessions) between the number of trials necessary to collect the last reward and the number of trials performed after obtention of the last reward failed to reach significance (rho <0.5, p>0.1 for all 3 monkeys). Still, at least at the beginning of the sessions, animals performed better than random agents, which implies that they were willing to mobilize cognitive resources to overcome the cognitive challenges, even partially.

**Figure 5:**
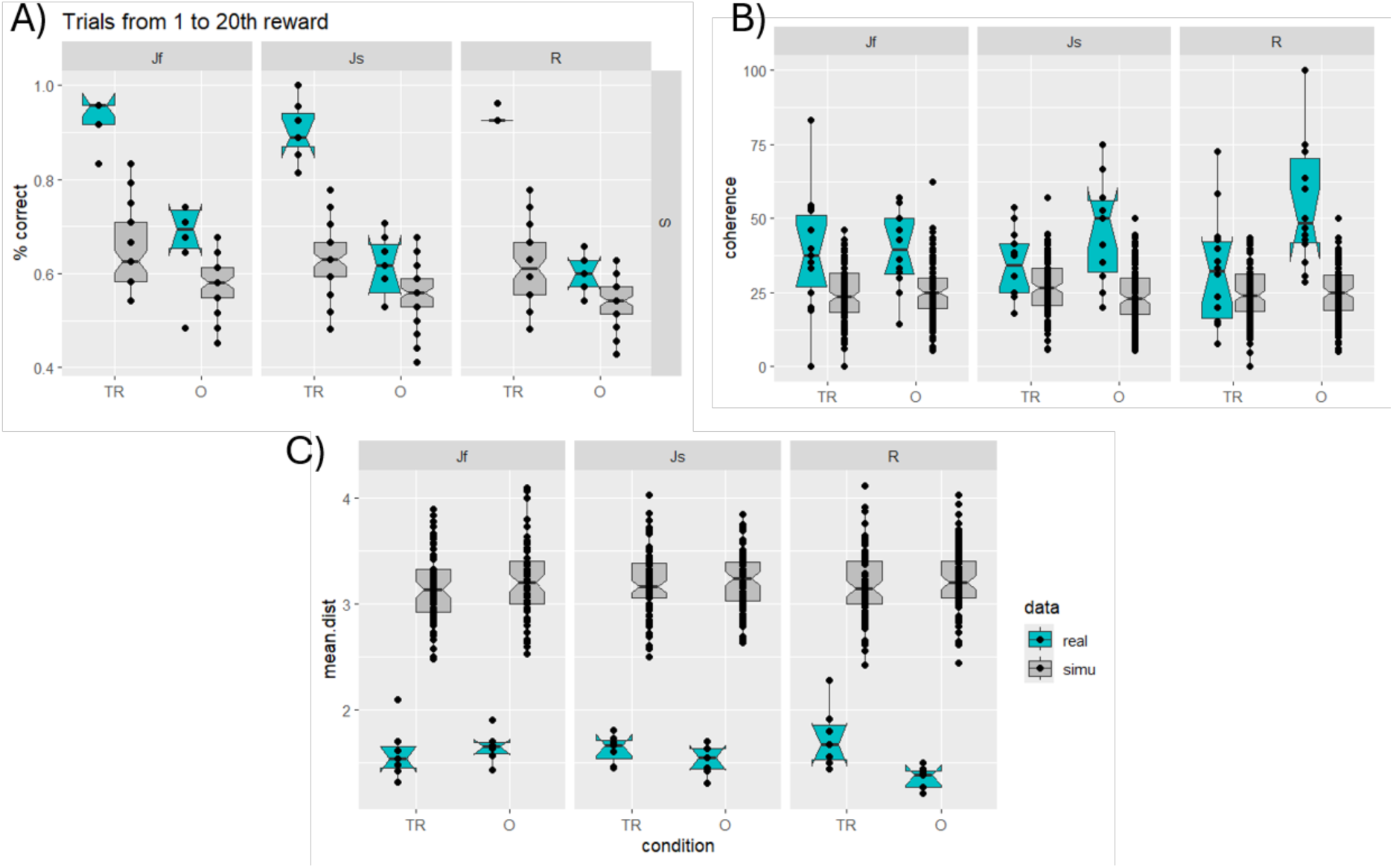
Performance (**A**), mean distance (**B**) and coherence measures (**C**) of monkeys and random agents in TR and O conditions for trials up to the obtention of the 20^th^ reward.

#### Spatial pattern of the behavior

We next examined the spatial patterns of movements, i.e. the relative position of successive choices across trials. For both O and TR conditions, we reasoned that an optimal agent should minimize the cost and thus the total distance covered to collect all the rewards. In addition, in opaque condition, such a systematic strategy would limit the working memory load. Thus, to avoid revisits and minimize the total traveled distance, animals should select neighboring wells and progress systematically along horizontal or vertical lines. In that frame, we can make simple predictions regarding the spatial pattern of movement displayed by an optimal agent: the inter-trial distance should not exceed 1 and the direction of successive actions should be coherent at least in one of the two dimensions. We computed the average distance between consecutive trials, as well as the coherence between successive actions both in the horizontal (x axis) and the vertical dimension (y axis). As for the average performance, we used computer simulations to generate the behavior of a random agent, which we used as a reference for the measures in real data.

The distance was computed on each trial as the sum of the number of wells between two consecutive choices along the X and Y axes. Selecting an adjacent well on the horizontal or vertical axis corresponded to a distance of 1, whereas a diagonal move corresponded to a distance of 2 (Δx=1 and Δy=1). We used ANOVAs to examine the variations of mean distance as a function of two factors: the type of data (real vs simulated data) and condition (opaque vs transparent doors). As shown on figure 5B, the average distance between successive choices was shorter in real data compared to simulations using random agents. For monkeys Jf and Js, only the main effect of data type was significant, with both monkeys making shorter movements in the real data compared to the simulations (Jf: F(1) = 261.73, p < 2 × 10⁻¹⁶; Js: F(1) = 371.11, p < 2 × 10⁻¹⁶). The average distance was undistinguishable between O and TR conditions (p>0.05). For monkey R, there was a significant interaction between the factor data type and the factor condition (F(1) = 5.82, p = 0.017). Post-hoc analysis revealed that distances tended to be shorter in O condition compared to TR in the real data (diff = –0.38, 95% CI [–0.80, 0.03], p = 0.079, Tukey test), but not in the simulated data (p>0.05). Similar results were obtained when using full sessions or only the first 20 trials of each session.

Coherence was calculated as the percentage of consecutive movements made in the same direction on each axis, and the data of the two axis were pooled for simplicity (Figure 5C). As for the average distance, we used ANOVAs to examine the influence of two factors: data type (real vs simulated data) and condition (opaque vs transparent doors). For all 3 monkeys, there was a significant effect of the factor data type (p<0.001), with a greater coherence in real data compared to simulated data (random agents). In addition, the interaction between data type and condition was significant for monkey R (F(1)=8.7, p=0.004)). In this animal, the percentage of coherent moves was significantly higher in opaque compared to transparent condition for real data (post hoc Tukey test, p=0.8 10^-4^), but not for simulated data using random agents (post hoc Tukey test, p=0.9). The level of coherence was indistinguishable between O and TR conditions for monkeys Jf and Js. Similar results were obtained when using full sessions or only the first 20 trials of each session. Note that similar results were obtained using non-parametric Wilcoxon signed-rank tests (supplementary material: ‘Supplementary analysis 3’).

In short, when considering the pattern of movements, all 3 monkeys were more efficient than random simulations in that they displayed shorter average distances and more coherence between successive moves. For monkey Jf and Js, we could not detect any difference in movement pattern between opaque and transparent conditions. By contrast, monkey R displayed both shorter distances and increased coherence in opaque compared to transparent condition. This suggests that in opaque condition, its movements were globally more efficient (in terms of minimizing distances), even if this increased efficacy was not associated with a better performance in the task, compared to the other monkeys.

### 2) Influence of DCZ injection on behavior

We next examined the influence of LC inactivation by comparing the influence of DCZ injection on different behavioral measures, in monkeys that did receive DREADD injections in the LC (Jf and Js). Monkey R did not receive DREADD injections in the LC and was used as a control, to assess the influence of DCZ injection alone. The dose of DCZ (0.1 mg/kg) was chosen based on its known influence on LC activity in monkey Jf, ie a 15% decrease in firing lasting over 30 minutes (Perez *et al*, 2022). Since Js received the same DREADD injection, it seems reasonable to assume that the influence of DCZ injection on LC activity was similar in this animal. In line with our previous study (Perez *et al*, 2022), we did not observe any effect of DCZ injection on vigilance or on locomotor activity in any of the 3 monkeys.

#### Transparent condition

We first examined the influence of DCZ injections in transparent condition, which was cognitively less demanding in that it did not require working memory. We compared monkeys’ success rate, distance and coherence values between saline and DCZ sessions (Figure 6).

**Figure 6.**
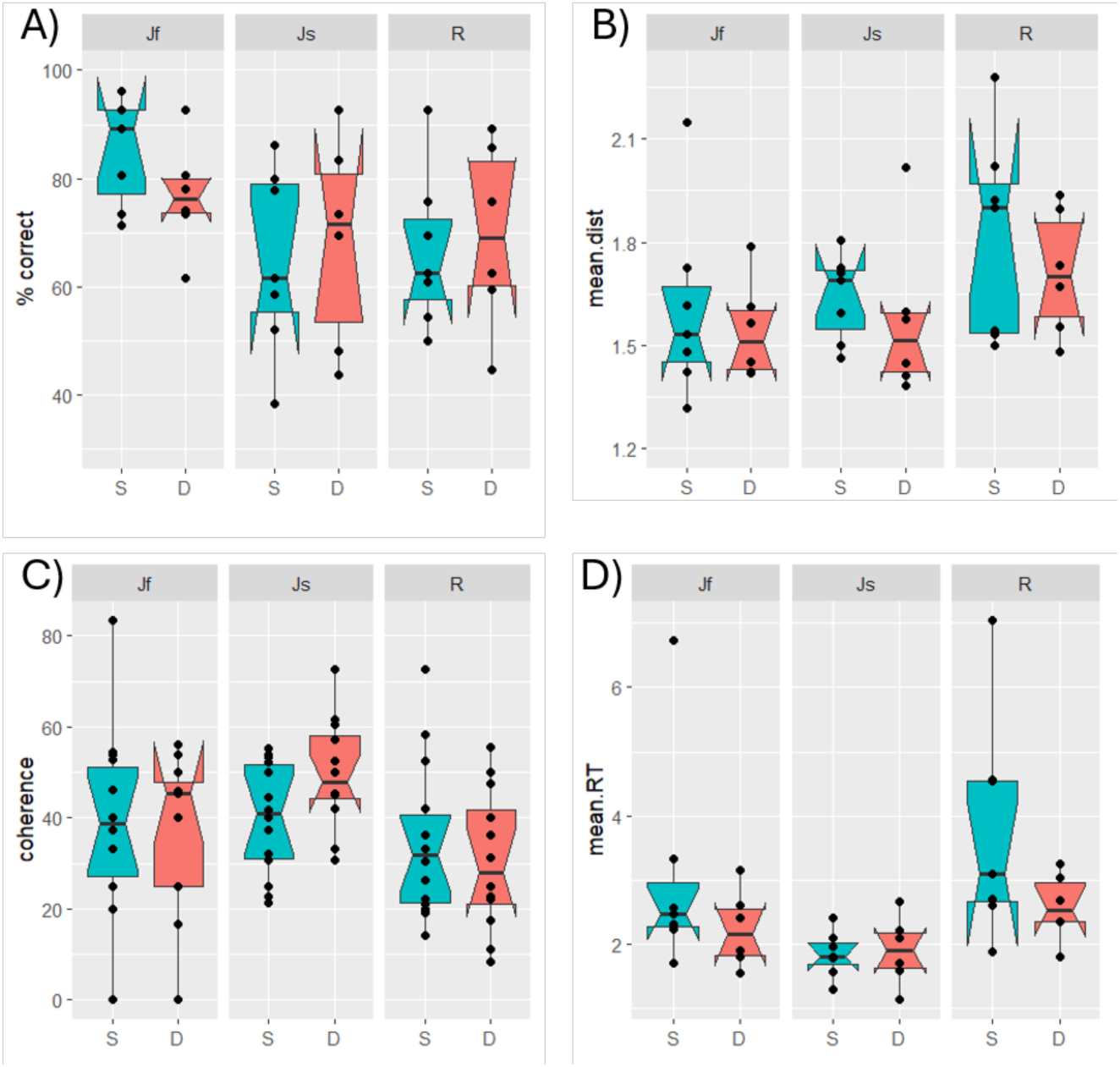
: Mean performance (A), mean distance (B), percentage of coherent movements (C) and mean RTs (D) in Transparent saline and DCZ conditions

In transparent condition, DCZ injections did not affect the mean performance (T-tests, Jf: t(10.31)=1.50, p=0.16; Js: t(10.19)=-0.35, p=0.74; R: t(9.88)=-0.35, p=0.73), nor the mean distance (T-tests, Jf: t(9.37)=0.53, p=0.61; Js: t(7.41)=0.64, p=0.54; R: t(10.07)=0.76, p=0.47) or the percentage of coherent movements (T-tests, Jf: t(48)=0.31, p=0.75; Js: t(41)=-1, p=0.3; R: t(50)=0.6, p=0.56) in any of the monkeys. Finally, we also examined the potential influence of DCZ on response times (RT), defined as the time difference between two consecutive choices. As for the other measures, mean RTs were not affected by DCZ in TR condition (T-tests, Jf: t(7.66) = 1.20, p = 0.27; Js: t(8.50) = -0.21, p = 0.84; R: t(7.23) = 1.72, p = 0.13). In short, the injection of DCZ did not seem to affect the behavior in transparent condition, where monkeys can use visual information to guide action selection. Note that similar results were obtained using non-parametric Wilcoxon signed-rank tests (supplementary material: ‘Supplementary analysis 4’).

#### Opaque condition

In opaque condition, which is more cognitively challenging in that in requires working memory, DCZ injection induced a decrease in performance for both monkeys injected with DREADDs in the LC (Figure 7A). Indeed, mean performance was significantly lower in DCZ sessions compared to Saline sessions (T-tests, Jf: t(9.23)=2.57, p=0.030; Js: t(11.98)=2.51, p=0.027). Note that even if the average performance decreased in DCZ sessions for these 2 monkeys, the number of rewards collected by session was not affected by the treatment (t.test, p>0.1 for both monkeys). Critically, DCZ did not affect performance in monkey R, who had not received DREADD injections t(10.55)=0.86, p=0.41). The treatment did not affect the number of rewards collected either (t(6.4)=1.2; p=0.2). Note that even if the performance of monkey R was relatively low compared to that of the other monkeys, it remained better than expected by random agent while rewards were available (see above, figure 5A). In addition, the number of trials performed after obtaining the last reward (by definition incorrect trials) was far from the maximum, since monkey R almost always quit the task long before the maximum session duration (5 minutes). Thus, even if in monkey R performance was low in saline condition, DCZ could still have decreased it by increasing the number of erroneous trials, before of after the obtention of the last reward. In summary, DCZ injection induced a decrease in performance in the 2 monkeys injected with inhibitory DREADDs in the LC, but not in the control monkey. Since DCZ did not affect the number of rewards obtained, the decrease in performance could only be related to an increase in incorrect trials. Indeed, monkeys Jf and Js performed overall more trials under DCZ (T-tests, Jf: t(11.01)= -2.50, p=0.030; Js: t(8.21)=-2.65, p=0.028) which was not the case of monkey R (T-test, t(12.72)=-0.95, p=0.36). Note that similar results were obtained using non-parametric Wilcoxon signed-rank tests (supplementary material: ‘Supplementary analysis 5’).

**Figure 7:**
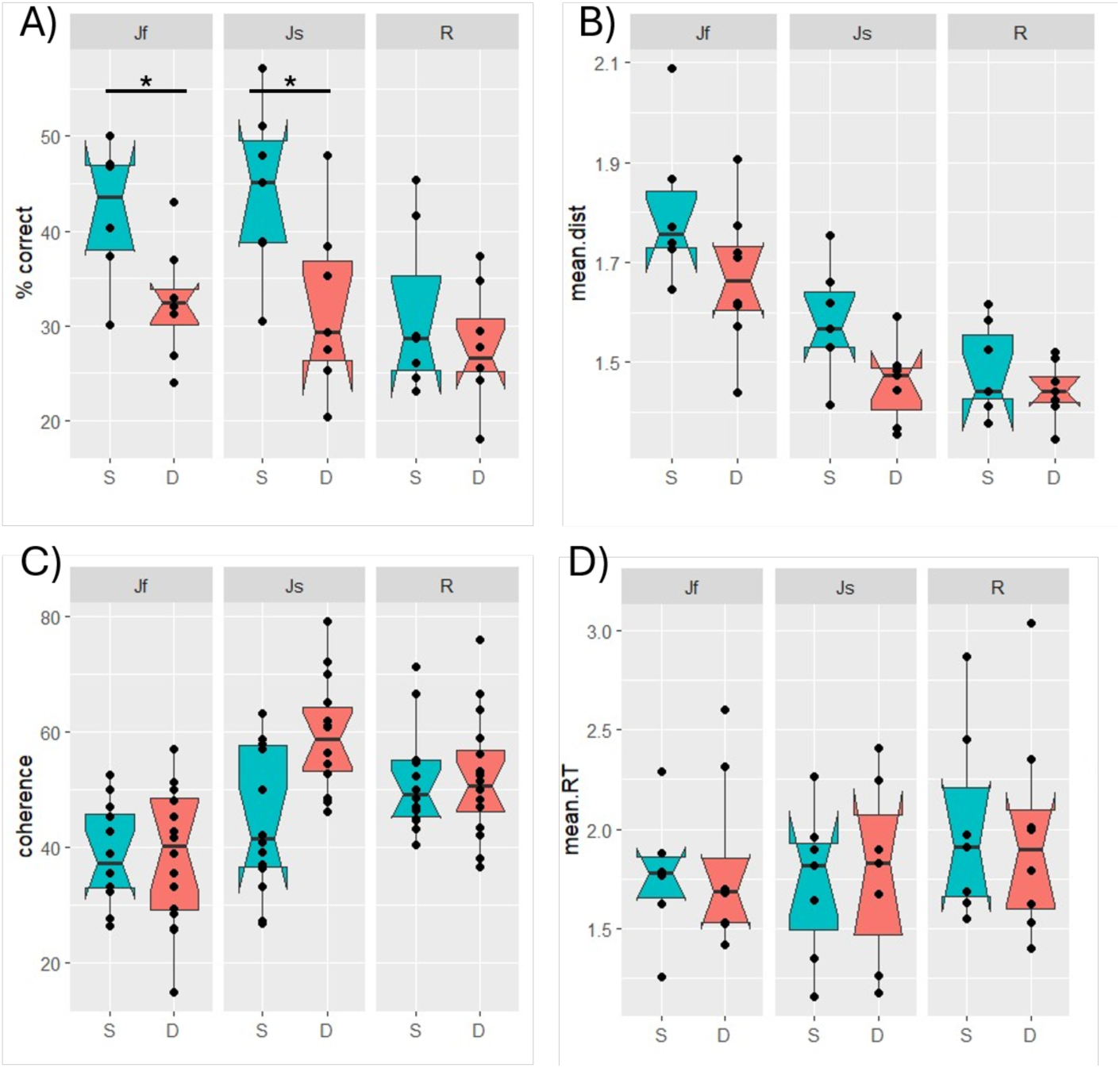
Mean performance (A), mean distance (B), percentage of coherent movements (C) and mean RTs (D) in Opaque saline and DCZ conditions

#### Dynamics of the effect

We first examined the influence of DCZ on the performance at the beginning of the sessions by only including trials up to the obtention of the 20th reward. The influence of DCZ on performance in the early part of the session did not reach significance for any of the 3 monkeys (T-test, Jf: t(7.4)=1.2, p=0.2; Js: t(8.5)=0.3, p=0.7; R: t(13)=0.4, p=0.7). We then investigated the influence of DCZ injection on the number of trials performed by the monkeys to retrieve the last reward of the session (note that this number could be different than 25). In both DREADD monkeys, the positive influence of DCZ on the number of trials to collect the last reward did not reach significance (T-tests, Jf: t(9.7)=1.2, p=0.26; Js: t(10.54)=-2.02, p=0.070). It was also the case in monkey R (T-test, t(12.70)=0.41, p=0.69). Because of the variability across sessions in the total number of rewards and the corresponding trial number, we ran a complementary analysis to examine the influence of DCZ on ability to collect a fixed number of rewards (Figure 8). We selected 23 rewards because it allowed us to include almost all the sessions (only 4 were excluded because monkeys collected less than 23 rewards: 1 of Jf, 2 for Js and 1 of R). DCZ induced an increase in the number of trials performed before the 23^rd^ reward for DREADD-injected monkeys. The effect reached significance for monkey Jf (T-tests, Jf: t(10.79) = -2.31, p = 0.042), but only approached significance for monkey Js (t(8.89) = 2.04, p = 0.072). No such effect could be observed for monkey R (T-test, t (11.61) = 0.89, p = 0.39). Thus, the influence DCZ injections in DREADD-injected monkeys did not appear at the very beginning of the sessions but it seemed to impair their ability to collect rewards in the later part of the session. No such effect could be observed in the control monkey which did not receive DREADDs in the LC.

**Figure 8:**
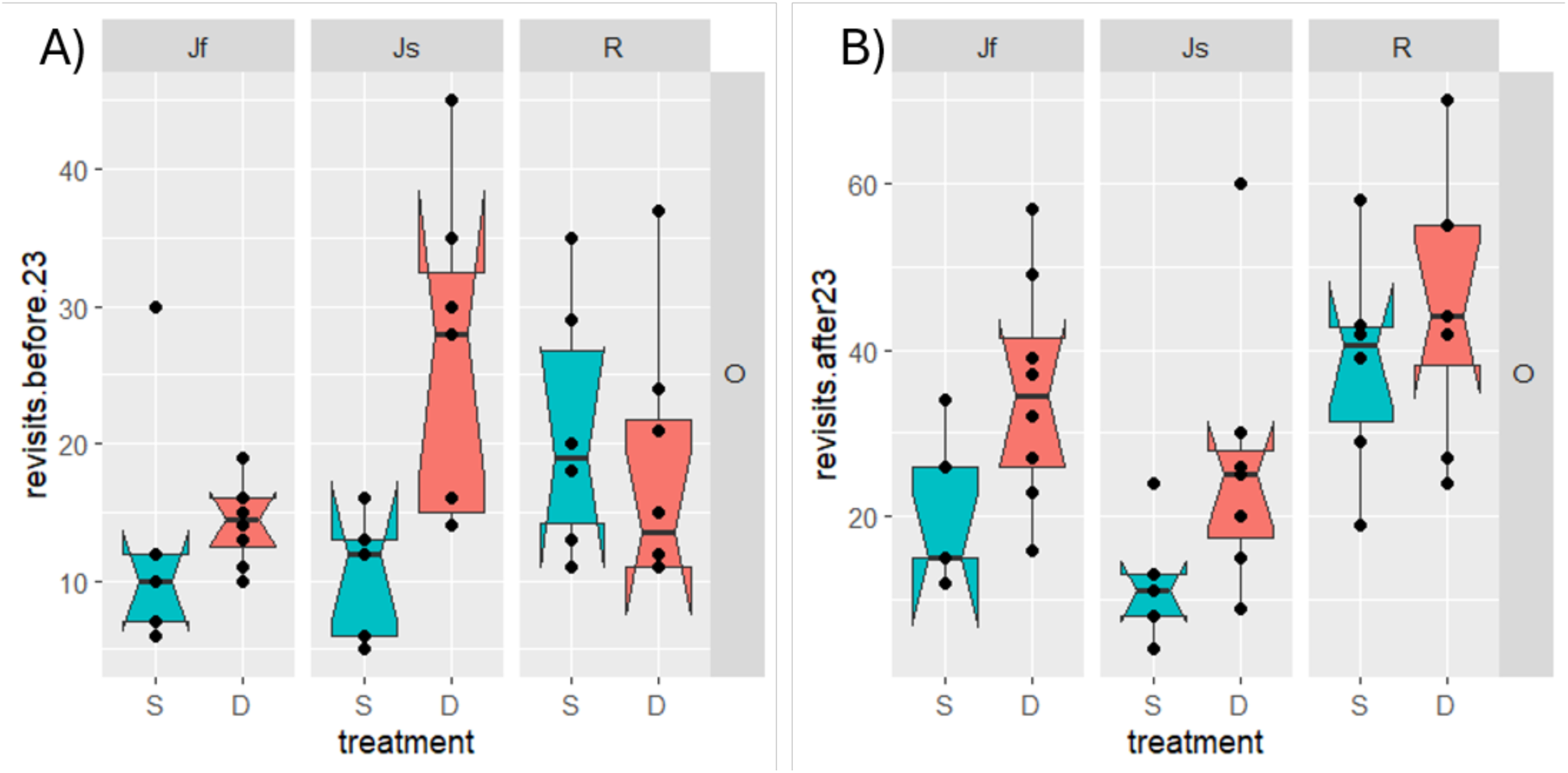
number of erroneous trials before (A) and after (B) retrieving the 23rd reward.

We also examined the influence of DCZ on the number of trials after the last reward, which, by definition are all erroneous. The influence of DCZ on this number only reached significance for monkey Js (T-test, t(7.2)=3.1, p=0.01) but neither for monkey Jf (T-test, t(10.9)=1.9, p=0.08) nor for monkey R (T-test, t(12.6)=1.9, p=0.07). Once again, in order to evaluate the dynamics of performance using a matched number of rewards, we examined the influence of DCZ on the number of erroneous trials performed after the 23rd reward (Figure 8B). The number of trials performed after the 23^rd^ reward increased under DCZ for monkey Js (T-test, t(10.79)=-2.31, p=0.042) and Jf only showed a non-significant trend (T-test, t(8.89)=-2.04, p=0.072). Monkey R showed no significant difference (T-test, t(11.60)=-0.89, p=0.39). In summary, in both monkeys injected with inhibitory DREADDs in the LC, DCZ injections increased the number of erroneous trials. This increase in erroneous responses was observed not only when monkeys were still collecting rewards (especially for monkey Jf), but also at the end of the sessions, when all (or virtually all) rewards had been collected already (especially for monkey Js). Note that similar results were obtained using non-parametric Wilcoxon signed-rank tests (supplementary material: ‘Supplementary analysis 6’). As it was the case in saline treatment, the correlation between the number of trials required to obtain the last reward and the number of trials after the obtention of the last reward failed to reach significance in all 3 monkeys, confirming the relative independence of these two measures.

#### Spatial pattern

We also measured the behavioral patterns under DCZ and compared them to those calculated under saline. As shown on Figure 7B, even though there is a tendency to a reduction of distance under DCZ, the effect was significant only in monkey Js (T-tests, t(11.1)=2.42, p=0.03). The effect did not reach significance for the other monkey injected with DREADDs (Jf: t(10.29)=1.71, p=0.12), nor for the control monkey (R: t(9.64)=-1.05, p=0.32). As shown on Figure 7C, the average percentage of coherence increased in DCZ sessions for monkey Js (t(3.4)=24.5, p=p=0.002). We did not observe any effect of DCZ on the percentage of coherence for the other monkey injected with DREADDs (Jf, t(25.9)=0.08, p=0.9) or for the control monkey (R, t(27.9)=0.04, p=0.97). Note that similar results were found when the analyses were run on the first part of the sessions (First 20 trials, see supplementary materials). As in transparent condition, we examined the potential influence of DCZ injection on response times (temporal delay between 2 consecutive trials). Once again, we found no effect of the drug on mean RTs (T-tests, Jf: t(11.90) = -0.20, p = 0.84; Js: t(11.57) = -0.26, p = 0.80; R: t(12.96) = 0.16, p = 0.87). Note that similar results were obtained using non-parametric Wilcoxon signed-rank tests (supplementary material: ‘Supplementary analysis 7’).

## Discussion

In summary, we trained monkeys in a task known to engage working memory and require the integrity of the DLPFC. In absence of treatment (saline conditions), all 3 monkeys could perform the task both in control (transparent doors) and ‘difficult’ (opaque doors) conditions, with only the later requiring working memory. Performance decreased in Opaque condition, as manifested both by a greater number of trials required to obtain the last reward and by a greater number of trials performed after the last reward was collected. But all monkeys always performed better than random agents. Since monkeys displayed similar exploration patterns when doors were transparent and opaques, they presumably mobilized additional cognitive resources in the later condition, to engage working memory and avoid revisits. Critically, decreasing LC activity with inhibitory DREADDs induced a decrease in performance in opaque condition, but not in transparent condition, which indicates that LC activity plays a critical role in enabling the mobilization of cognitive resources to support working memory performance.

The task that we chose was simple and monkeys learned it quickly. Compared to more complex tasks used in laboratory settings to study working memory, this one involves a spontaneous behavior (searching rewards in the wells, by turning the doors) and it virtually did not require any training (Miller, Erickson et al. 1996, Wittig, Morgan et al. 2016). After a first session where monkeys were exposed to the hole-board, with a reward in each well, they spontaneously displaced the doors to access the reward. Critically, once the animals had understood that each well contained only one reward such that all subsequent visits after the first one would not yield anything, they spontaneously avoided revisits (Figure 2). This is in line with the idea that, all other things being equal, animals tend to maximize the ratio between benefits (here the number of rewards obtained) and the costs (the number of actions performed). In opaque condition, given the lack of visual information, animals had the same goal but they needed to memorize past actions to avoid revisits, which represented a clear cognitive challenge. Indeed, error rates increased, even if they remained below that of random agent simulations. Thus, as in past studies using the same task, monkeys spontaneously engaged cognitive resources to avoid revisits in the absence of visual feedback, at least in part by engaging working memory (Collin, Cowey et al. 1982, Passingham 1985).

The optimal strategy in this task does not only consist in avoiding revisits but also in minimizing the total distance covered to collect all 25 rewards. Practically, animals should progress along straight lines (horizontal or vertical) and select immediately neighboring wells. We showed that monkeys were far less organized than an optimal agent, since their average distance between successive choices was greater than one and the coherence in the direction of successive actions was lower than 100%. This is in line with previous studies (Collin, Cowey et al. 1982, Passingham 1985, Gandaux, Sallet et al. 2025). But their behavior was still significantly more organized than expected from a random agent (figure 5). Previous studies showed that the spatio-temporal organization of behavior in this task, even if not optimal, could be significantly impaired by DLPFC lesions (Collin, Cowey et al. 1982, Passingham 1985). This was also in line with early studies in humans showing the role of the PFC in spatial working memory and strategic planning (Owen, Downes et al. 1990). Note that for 2 of the 3 monkeys, the level of organization (as measured using distance and coherence) was similar between transparent and opaque conditions. And even if the third animal tended to show a better organization of behavior in Opaque compared to Transparent condition, this animal was showing a slightly poorer average performance compared to the other two, such that this change in behavioral organization did not seem to enhance performance. Last, a similar pattern was reported in a recent study using the same task, in that the fraction of short distance moves was similar between transparent and opaque conditions (Gandaux, Sallet et al. 2025). Thus, it seems unlikely that monkeys adjust to the opaque condition by adopting a distinct exploration strategy. Rather, they seem to be using the same general strategy, which results in decreased distance and increased coherence between successive actions compared to a random agent, even in Transparent condition. In the absence of visual information in Opaque condition, monkeys presumably engage working memory to keep track of past actions and limit revisits (See also Passingham, 1985). This enables them to perform better than chance level and collect as many rewards as in transparent condition, even if the difficulty is such that they require more trials. Altogether, this is compatible with the idea that in opaque condition, monkeys engage more cognitive resources to perform the task using working memory, which we interpret in terms of increased cognitive effort.

We used a targeted pharmacogenetic approach to decrease the activity of LC neurons while monkeys were performing the task, both in Transparent and Opaque conditions. This allowed us to evaluate the contribution of the LC as a whole, rather than the influence of NA on a subset of brain regions, as a function of receptor subtypes, as it is the case with pharmacology (Arnsten 2000, Berridge and Waterhouse 2003, Poe, Foote et al. 2020, Grimm, Duss et al. 2024, Xia, Maheu et al. 2026). As in previous studies using the same dose of DCZ to activate the DREADDs, DCZ alone had no effect on behavior (Nagai, Miyakawa et al. 2020, Gandaux, Sallet et al. 2025). Using an expression vector carrying a Tyrosine Hydroxylase promoter, we ensured that only LC neurons expressed the inhibitory DREADD hM4Di (Perez et al, 2020). Using neurophysiological recordings, we had shown that injecting DCZ (0.1 mg/kg) in one of these monkeys induced a rapid (within minutes) and reversible decrease of LC activity (to about 85% of pre-injection firing rate), which lasted at least one hour (Perez, Chavret-Reculon et al. 2022). This is clearly enough for our purpose here, given that individual sessions in this task started 30 minutes after DCZ injection and never exceeded 5 minutes. Thus, we could be relatively confident that during these experiments, the injection of DCZ activated the inhibitory DREADDs in the LC and induced a moderate (15%) but reliable decrease in LC activity for the entire duration of the experiment.

This moderate inhibition did not induce any visible decrease in vigilance or arousal, but it clearly affected behavior in the task. Indeed, the treatment induced an increase in revisits in opaque condition, but not in transparent condition. This is in line with previous pharmacological studies showing that a limited decrease in LC activity could induce task-related behavioral effects without significantly altering vigilance or arousal (Riekkinen, Kejonen et al. 1998, Jahn, Gilardeau et al. 2018, Borderies, Bornert et al. 2020, Dubois, Habicht et al. 2021). This is also in line with our previous study showing that at that dose (0.1 mg/kg), DCZ induced a 15% decrease in LC activity but no change in vigilance or locomotor activity (Perez, Chavret-Reculon et al. 2022). In addition, the deficit in performance was subtle in that it did not prevent the animals from obtaining as many rewards as they would under saline treatment, but it only increased the number of trials necessary to obtain it. Finally, the treatment did not affect response times. Thus, the treatment did not affect basic sensory and motor functions, and it did not influence the global level of motivation of the monkeys. Thus, the decreased performance induced by LC inhibition in opaque conditions is likely related to a deficit in working memory, as showed earlier in the same task with DLPFC lesions (Collin, Cowey et al. 1982, Passingham 1985). Note, however, that the effects of a moderate and reversible LC inhibition were more subtle than the effect of DLPFC lesions described in previous studies. For example, DLPFC lesions induced performance deficits that could be observed from the very onset of the task: in Collin et al (1982), DLPFC lesions induced a clear increase in the number of trials required to obtain 20 rewards, which is not the case here with LC inactivation. In addition, moderate LC inhibition did not induce a decrease in behavioral organization, as it could be observed after DLPFC lesions (Collin, Cowey et al. 1982, Passingham 1985). Here, even if the effects were small, they were fairly specific and resulted in an increase in number of erroneous trials. Given the task requirements (selecting wells that have not been visited before), it seems likely that LC inhibition alters the ability to remember previously visited wells and plan accordingly, to select rewarded locations. This is compatible with an interpretation in terms of working memory impairment. But the inhibition of LC neurons also induced an increase in the number of trials performed after the last reward had been collected. Since this effect was only observed in Opaque condition, it probably does not reflect a simple increase in perseveration or a compulsive behavior. Rather, LC inhibition could affect the ability to keep track of the progression through the task, and to disengage when monkeys know (or believe) that there is no reward left. In transparent condition, this can be achieved by simple visual control and LC inhibition has no effect. In Opaque condition, monkeys presumably need to use working memory to keep track of their past successes and failures to infer that, at some point, all rewards have been collected and they should stop performing the task. In that frame, LC inhibition would alter the ability to keep track of past actions and plan accordingly, both to select rewards when there are still some, and to stop performing the task once all rewards have been retrieved. Again, this is compatible with an interpretation in terms of working memory.

Interestingly, a similar increase in the number of trials performed was recently described in monkeys performing the same task, after specific activation of the MCC-DLPFC pathway, and interpreted in terms of increased engagement (Gandaux, Sallet et al. 2025). Given the strong functional overlap between LC and MCC, often associated with cognitive control, it seems surprising that LC inhibition and MCC-DLPFC activation led to such similar behavioral patterns (Aston-Jones and Cohen 2005, van der Wel and van Steenbergen 2018, Muller, Mars et al. 2019). But critically, LC inhibition primarily led to a decrease in performance, which was not the case for MCC activation. In that frame, we could interpret the increased number of trials in the present study as a secondary increase in engagement, to compensate for the decreased performance induced by the LC inhibition. Critically, both in Gandaux et al (2025) and in the present study, this increased number of trials performed is not instrumental, in that it does not increase the amount of rewards obtained. It even resulted in a clear decrease in the benefits/cost ratio, which implies a deficit of cognitive (or metacognitive) processes controlling resources allocation. In summary, whether monkeys performed more trials because they have a deficit in identifying the point at which they should stop (because reward rate is too low) or whether they allocate excessive resources in the task (to perform trials when virtually all rewards have been collected), it points to an alteration of executive control functions.

In sum, the pattern of behavioral changes induced by reversible LC inhibition seems to be related to an alteration of executive functions, rather than a global effect in terms of motivation or vigilance. This is in line with the known positive influence of NA on working memory, through an activation of local alpha-2 receptors (Arnsten 2000). The current work demonstrates that this positive influence on working memory can be obtained by manipulating LC activity as a whole, and not only by manipulating local NA action in the PFC. We believe, however, that the role of the LC extends beyond working memory. Indeed, systemic manipulations of the NA system have been proven to influence other cognitive functions such as attention and inhibitory control (Witte and Marrocco 1997, Chamberlain, Müller et al. 2006, Reynaud, Froesel et al. 2019). It is thus tempting to interpret the deficit associated with LC/NA alteration in terms of cognitive control/mental effort (Westbrook and Braver 2015, Shenhav, Musslick et al. 2017, Unsworth and Robison 2017, Unsworth, Miller et al. 2022). Indeed, our previous studies had already identified a strong correlation between the activation of LC neurons and the allocation of both physical and cognitive effort, in situations that did not involve any kind of working memory (Varazzani, San-Galli et al. 2015, Bornert and Bouret 2021). We had also shown that noradrenaline was causally involved in physical effort (Borderies, Bornert et al. 2020). Thus, the present work extends the implication of the LC/NA system to cognitive effort, such that the activation of LC and the corresponding release of NA in target areas would allow the increase in resources underlying performance both in the physical and in the cognitive domain.

This idea is in line with numerous studies showing the strong relation between autonomic arousal (and in particular pupil dilation) and cognitive effort (Kahneman and Beatty 1966, Peters, Godaert et al. 1998, Howells, Stein et al. 2010, Zenon 2014, Kuipers, Richter et al. 2017, van der Wel and van Steenbergen 2018). Several authors interpret the relation between effort and pupil in terms of LC activity (Unsworth and Robison 2017, van der Wel and van Steenbergen 2018, Unsworth, Robison et al. 2025). Indeed, the firing of LC neurons is strongly related to autonomic arousal (Abercrombie and Jacobs 1987, Aston-Jones, Rajkowski et al. 1996). But inferring the role of the LC/ NA system remains subject to caution, given that autonomic arousal is related to a vast number of brain regions, both cortical and subcortical (Saper 2002, Barbas and Saha 2003, Critchley 2009, Joshi, Li et al. 2016, Reimer, McGinley et al. 2016). Thus, the present study provides a critical complement to these studies based on indirect measures, to establish the the critical contribution of the LC/NA system to cognitive effort. Finally, the mechanisms underlying this function remain very hypothetical, since LC activity influences numerous cortical and subcortical regions, targeting various receptor subtypes and various neural populations (Robbins and Arnsten 2009, Poe, Foote et al. 2020). Recent studies also indicate that LC activation could act on behavior through its influence on glial cells, which could regulate the amount of energy allocated to the neurons (Grimm, Duss et al. 2024, Lefton, Wu et al. 2025).

In conclusion, this work indicates that the LC plays a significant causal role in cognitive effort, in addition to its known role in physical effort (Borderies, Bornert et al. 2020). This is in line with recent studies showing the strong relation between physical and cognitive effort both in healthy subjects and in depressed patients (Vinckier, Jaffre et al. 2022, Bustamante, Oshinowo et al. 2023)). Further research would be necessary to understand the underlying cellular mechanisms, and also the clinical implications.

## References

Abercrombie, E. D. and B. L. Jacobs (1987). “Single-unit response of noradrenergic neurons in the locus coeruleus of freely moving cats. I. Acutely presented stressful and nonstressful stimuli.” The Journal of neuroscience : the official journal of the Society for Neuroscience 7(9): 2837–2843.

Alnaes, D., M. H. Sneve, T. Espeseth, T. Endestad, S. H. van de Pavert and B. Laeng (2014). “Pupil size signals mental effort deployed during multiple object tracking and predicts brain activity in the dorsal attention network and the locus coeruleus.” J Vis 14(4).

Arnsten, A. F. T. (2000). “Through the looking glass: differential noradenergic modulation of prefrontal cortical function.” Neural plasticity 7(1-2): 133–146.

Aston-Jones, G. and J. D. Cohen (2005). “An integrative theory of locus coeruleus-norepinephrine function: adaptive gain and optimal performance.” Annual Review of Neuroscience 28: 403–450.

Aston-Jones, G., J. Rajkowski, P. Kubiak, R. J. Valentino and M. T. Shipley (1996). “Role of the locus coeruleus in emotional activation.” Progress in brain research 107: 379–402.

Bacon, T. J., A. E. Pickering and J. R. Mellor (2020). “Noradrenaline Release from Locus Coeruleus Terminals in the Hippocampus Enhances Excitation-Spike Coupling in CA1 Pyramidal Neurons Via beta-Adrenoceptors.” Cereb Cortex 30(12): 6135–6151.

Barbas, H. and S. Saha (2003). “Serial pathways from primate prefrontal cortex to autonomic areas may influence emotional expression.” BMC neuroscience 4: 25.

Berridge, C. W. and B. D. Waterhouse (2003). “The locus coeruleus-noradrenergic system: modulation of behavioral state and state-dependent cognitive processes.” Brain research Brain research reviews 42(1): 33–84.

Borderies, N., P. Bornert, S. Gilardeau and S. Bouret (2020). “Pharmacological evidence for the implication of noradrenaline in effort.” PLoS Biol 18(10): e3000793.

Bornert, P. and S. Bouret (2021). “Locus coeruleus neurons encode the subjective difficulty of triggering and executing actions.” PLoS Biol 19(12): e3001487.

Botvinick, M. M., T. S. Braver, D. M. Barch, C. S. Carter and J. D. Cohen (2001). “Conflict monitoring and cognitive control.” Psychological review 108(3): 624.

Bustamante, L. A., T. Oshinowo, J. R. Lee, E. Tong, A. R. Burton, A. Shenhav, J. D. Cohen and N. D. Daw (2023). “Effort Foraging Task reveals positive correlation between individual differences in the cost of cognitive and physical effort in humans.” Proc Natl Acad Sci U S A 120(50): e2221510120.

Cerpa, J. C., A. Piccin, M. Dehove, M. Lavigne, E. J. Kremer, M. Wolff, S. L. Parkes and E. Coutureau (2023). “Inhibition of noradrenergic signalling in rodent orbitofrontal cortex impairs the updating of goal-directed actions.” Elife 12.

Chamberlain, S. R., A. Hampshire, U. Muller, K. Rubia, N. Del Campo, K. Craig, R. Regenthal, J. Suckling, J. P. Roiser, J. E. Grant, E. T. Bullmore, T. W. Robbins and B. J. Sahakian (2009). “Atomoxetine modulates right inferior frontal activation during inhibitory control: a pharmacological functional magnetic resonance imaging study.” Biol Psychiatry 65(7): 550–555.

Chamberlain, S. R., U. Müller, A. D. Blackwell, L. Clark, T. W. Robbins and B. J. Sahakian (2006). “Neurochemical modulation of response inhibition and probabilistic learning in humans.” Science (New York, NY) 311(5762): 861–863.

Collin, N. G., A. Cowey, R. Latto and C. Marzi (1982). “The role of frontal eye-fields and superior colliculi in visual search and non-visual search in rhesus monkeys.” Behav Brain Res 4(2): 177–193.

Critchley, H. D. (2009). “Psychophysiology of neural, cognitive and affective integration: fMRI and autonomic indicants.” International journal of psychophysiology : official journal of the International Organization of Psychophysiology: 1–7.

de Gee, J. W., K. Tsetsos, L. Schwabe, A. E. Urai, D. McCormick, M. J. McGinley and T. H. Donner (2020). “Pupil-linked phasic arousal predicts a reduction of choice bias across species and decision domains.” Elife 9.

Dubois, M., J. Habicht, J. Michely, R. Moran, R. J. Dolan and T. U. Hauser (2021). “Human complex exploration strategies are enriched by noradrenaline-modulated heuristics.” Elife 10.

Einhäuser, W., J. Stout, C. Koch and O. L. Carter (2008). “Pupil dilation reflects perceptual selection and predicts subsequent stability in perceptual rivalry.” Proceedings of the National Academy of Sciences of the United States of America 105(5): 1704–1709.

Filipowicz, A. L., C. M. Glaze, J. W. Kable and J. I. Gold (2020). “Pupil diameter encodes the idiosyncratic, cognitive complexity of belief updating.” Elife 9.

Gandaux, C., J. Sallet, C. Amiez, D. Autran-Clavagnier, V. Morel-Latour, C. Goussi-Denjean, V. Fontanier, P. Misery, C. Lamy, F. Lamberton, M. Lavigne, E. J. Kremer, C. R. E. Wilson and E. Procyk (2025). “Motivational control is implemented by a cingulo-prefrontal network.” Curr Biol 35(22): 5618–5625 e5614.

Grimm, C., S. N. Duss, M. Privitera, B. R. Munn, N. Karalis, S. Frassle, M. Wilhelm, T. Patriarchi, D. Razansky, N. Wenderoth, J. M. Shine, J. Bohacek and V. Zerbi (2024). “Tonic and burst-like locus coeruleus stimulation distinctly shift network activity across the cortical hierarchy.” Nat Neurosci 27(11): 2167–2177.

Hernaus, D., M. M. Casales Santa, J. S. Offermann and T. Van Amelsvoort (2017). “Noradrenaline transporter blockade increases fronto-parietal functional connectivity relevant for working memory.” Eur Neuropsychopharmacol 27(4): 399–410.

Hirschberg, S., Y. Li, A. Randall, E. J. Kremer and A. E. Pickering (2017). “Functional dichotomy in spinal- vs prefrontal-projecting locus coeruleus modules splits descending noradrenergic analgesia from ascending aversion and anxiety in rats.” Elife 6.

Howells, F. M., D. J. Stein and V. A. Russell (2010). “Perceived mental effort correlates with changes in tonic arousal during attentional tasks.” Behavioral and Brain Functions 6(1): 39.

Jacobs, B. L., E. D. Abercrombie, C. A. Fornal, E. S. Levine, D. A. Morilak and I. L. Stafford (1991). “Single-unit and physiological analyses of brain norepinephrine function in behaving animals.” Progress in brain research 88: 159–165.

Jahn, C. I., S. Gilardeau, C. Varazzani, B. Blain, J. Sallet, M. E. Walton and S. Bouret (2018). “Dual contributions of noradrenaline to behavioural flexibility and motivation.” 1–16.

Joshi, S., Y. Li, R. M. Kalwani and J. I. Gold (2016). “Relationships between Pupil Diameter and Neuronal Activity in the Locus Coeruleus, Colliculi, and Cingulate Cortex.” Neuron 89(1): 221–234.

Kahneman, D. and J. Beatty (1966). “Pupil diameter and load on memory.” Science (New York, NY).

Kuipers, M., M. Richter, D. Scheepers, M. A. Immink, E. Sjak-Shie and H. van Steenbergen (2017). “How effortful is cognitive control? Insights from a novel method measuring single-trial evoked beta-adrenergic cardiac reactivity.” Int J Psychophysiol 119: 87–92.

Lefton, K. B., Y. Wu, Y. Dai, T. Okuda, Y. Zhang, A. Yen, G. M. Rurak, S. Walsh, R. Manno, B. E. Myagmar, J. D. Dougherty, V. K. Samineni, P. C. Simpson and T. Papouin (2025). “Norepinephrine signals through astrocytes to modulate synapses.” Science 388(6748): 776–783.

Lisi, M., M. Bonato and M. Zorzi (2015). “Pupil dilation reveals top–down attentional load during spatial monitoring.” Biological psychology 112(C): 39–45.

Miller, E. K., C. A. Erickson and R. Desimone (1996). “Neural mechanisms of visual working memory in prefrontal cortex of the macaque.” J Neurosci 16(16): 5154–5167.

Muller, T. H., R. B. Mars, T. E. J. Behrens and J. X. O’Reilly (2019). “Control of entropy in neural models of environmental state.” Elife 8.

Murphy, P. R., R. G. O’Connell, M. O’Sullivan, I. H. Robertson and J. H. Balsters (2014). “Pupil diameter covaries with BOLD activity in human locus coeruleus.” Human Brain Mapping 35(8): 4140–4154.

Naccache, L., S. Dehaene, L. Cohen, M.-O. Habert, E. Guichart-Gomez, D. Galanaud and J.-C. Willer (2005). “Effortless control: executive attention and conscious feeling of mental effort are dissociable.” Neuropsychologia 43(9): 1318–1328.

Nagai, Y., N. Miyakawa, H. Takuwa, Y. Hori, K. Oyama, B. Ji, M. Takahashi, X. P. Huang, S. T. Slocum, J. F. DiBerto, Y. Xiong, T. Urushihata, T. Hirabayashi, A. Fujimoto, K. Mimura, J. G. English, J. Liu, K. I. Inoue, K. Kumata, C. Seki, M. Ono, M. Shimojo, M. R. Zhang, Y. Tomita, J. Nakahara, T. Suhara, M. Takada, M. Higuchi, J. Jin, B. L. Roth and T. Minamimoto (2020). “Deschloroclozapine, a potent and selective chemogenetic actuator enables rapid neuronal and behavioral modulations in mice and monkeys.” Nat Neurosci 23(9): 1157–1167.

Nassar, M. R., K. M. Rumsey, R. C. Wilson, K. Parikh, B. Heasly and J. I. Gold (2012). “Rational regulation of learning dynamics by pupil-linked arousal systems.” Nature Neuroscience 15(7): 1040–1046.

Negelspach, D., A. Alkozei, A. Huskey and W. D. S. Killgore (2025). “Effects of Locus Coeruleus Activation on n-Back Performance and Frontoparietal Activity.” J Cogn Neurosci 37(1): 97–109.

Owen, A. M., J. J. Downes, B. J. Sahakian, C. E. Polkey and T. W. Robbins (1990). “Planning and spatial working memory following frontal lobe lesions in man.” Neuropsychologia 28(10): 1021–1034.

Passingham, R. (1985). “Memory of monkeys (Macaca mulatta) with lesions in prefrontal cortex.” Behavioral neuroscience 99(1): 3.

Perez, P., E. Chavret-Reculon, P. Ravassard and S. Bouret (2022). “Using Inhibitory DREADDs to Silence LC Neurons in Monkeys.” Brain Sci 12(2).

Peters, M. L., G. L. Godaert, R. E. Ballieux, M. van Vliet, J. J. Willemsen, F. C. Sweep and C. J. Heijnen (1998). “Cardiovascular and endocrine responses to experimental stress: effects of mental effort and controllability.” Psychoneuroendocrinology 23(1): 1–17.

Poe, G. R., S. Foote, O. Eschenko, J. P. Johansen, S. Bouret, G. Aston-Jones, C. W. Harley, D. Manahan-Vaughan, D. Weinshenker, R. Valentino, C. Berridge, D. J. Chandler, B. Waterhouse and S. J. Sara (2020). “Locus coeruleus: a new look at the blue spot.” Nat Rev Neurosci.

Pribram, K. H. and D. McGuinness (1975). “Arousal, activation, and effort in the control of attention.” Psychol Rev 82(2): 116–149.

Rae, C. L., C. Nombela, P. V. Rodriguez, Z. Ye, L. E. Hughes, P. S. Jones, T. Ham, T. Rittman, I. Coyle-Gilchrist, R. Regenthal, B. J. Sahakian, R. A. Barker, T. W. Robbins and J. B. Rowe (2016). “Atomoxetine restores the response inhibition network in Parkinson’s disease.” Brain 139(Pt 8): 2235–2248.

Reimer, J., M. J. McGinley, Y. Liu, C. Rodenkirch, Q. Wang, D. A. McCormick and A. S. Tolias (2016). “Pupil fluctuations track rapid changes in adrenergic and cholinergic activity in cortex.” Nat Commun 7: 13289.

Reynaud, A. J., M. Froesel, C. Guedj, S. Ben Hadj Hassen, J. Cléry, M. Meunier, S. Ben Hamed and F. Hadj-Bouziane (2019). “Atomoxetine improves attentional orienting in a predictive context.” Neuropharmacology 150: 59–69.

Riekkinen, M., K. Kejonen, P. Jäkälä, H. Soininen and P. Riekkinen (1998). “Reduction of noradrenaline impairs attention and dopamine depletion slows responses in Parkinson’s disease.” The European journal of neuroscience 10(4): 1429–1435.

Robbins, T. W. and A. F. T. Arnsten (2009). “The neuropsychopharmacology of fronto-executive function: monoaminergic modulation.” Annual Review of Neuroscience 32(1): 267–287.

Rossetti, Z. L. and S. Carboni (2005). “Noradrenaline and dopamine elevations in the rat prefrontal cortex in spatial working memory.” The Journal of neuroscience : the official journal of the Society for Neuroscience 25(9): 2322–2329.

Saper, C. B. (2002). “The central autonomic nervous system: conscious visceral perception and autonomic pattern generation.” Annual Review of Neuroscience 25: 433–469.

Shenhav, A., S. Musslick, F. Lieder, W. Kool, T. L. Griffiths, J. D. Cohen and M. M. Botvinick (2017). “Toward a Rational and Mechanistic Account of Mental Effort.” Annu Rev Neurosci 40: 99–124.

Tomassini, A., F. H. Hezemans, R. Ye, K. A. Tsvetanov, N. Wolpe and J. B. Rowe (2022). “Prefrontal Cortical Connectivity Mediates Locus Coeruleus Noradrenergic Regulation of Inhibitory Control in Older Adults.” J Neurosci 42(16): 3484–3493.

Unsworth, N., A. L. Miller and S. Aghel (2022). “Effort Mobilization and Lapses of Sustained Attention.” Cogn Affect Behav Neurosci 22(1): 42–56.

Unsworth, N. and M. K. Robison (2017). “A locus coeruleus-norepinephrine account of individual differences in working memory capacity and attention control.” Psychon Bull Rev 24(4): 1282–1311.

Unsworth, N., M. K. Robison and A. L. Miller (2025). “Mobilizing effort to reduce lapses of sustained attention: examining the effects of content-free cues, feedback, and points.” Cogn Affect Behav Neurosci 25(3): 631–649.

van der Wel, P. and H. van Steenbergen (2018). “Pupil dilation as an index of effort in cognitive control tasks: A review.” 1–11.

Varazzani, C., A. San-Galli, S. Gilardeau and S. Bouret (2015). “Noradrenaline and Dopamine Neurons in the Reward/Effort Trade-Off: A Direct Electrophysiological Comparison in Behaving Monkeys.” The Journal of neuroscience : the official journal of the Society for Neuroscience 35(20): 7866–7877.

Vinckier, F., C. Jaffre, C. Gauthier, S. Smajda, P. Abdel-Ahad, R. Le Bouc, J. Daunizeau, M. Fefeu, N. Borderies, M. Plaze, R. Gaillard and M. Pessiglione (2022). “Elevated Effort Cost Identified by Computational Modeling as a Distinctive Feature Explaining Multiple Behaviors in Patients With Depression.” Biol Psychiatry Cogn Neurosci Neuroimaging 7(11): 1158–1169.

Westbrook, A. and T. S. Braver (2015). “Cognitive effort: A neuroeconomic approach.” Cognitive, affective & behavioral neuroscience 15(2): 395–415.

Witte, E. A. and R. T. Marrocco (1997). “Alteration of brain noradrenergic activity in rhesus monkeys affects the alerting component of covert orienting.” Psychopharmacology (Berl) 132(4): 315–323.

Wittig, J. H., Jr., B. Morgan, E. Masseau and B. J. Richmond (2016). “Humans and monkeys use different strategies to solve the same short-term memory tasks.” Learn Mem 23(11): 644–647.

Xia, H., M. Maheu, G. A. Kane and B. B. Scott (2026). “Regulation of the decision threshold by the locus coeruleus.” Neuropsychopharmacology.

Zenon, A. (2014). “Pupil size variations correlate with physical effort perception.” 1–8.

Zenon, A., M. Sidibe and E. Olivier (2015). “Disrupting the Supplementary Motor Area Makes Physical Effort Appear Less Effortful.” The Journal of neuroscience : the official journal of the Society for Neuroscience 35(23): 8737–8744.

Zerbi, V., A. Floriou-Servou, M. Markicevic, Y. Vermeiren, O. Sturman, M. Privitera, L. von Ziegler, K. D. Ferrari, B. Weber, P. P. De Deyn, N. Wenderoth and J. Bohacek (2019). “Rapid Reconfiguration of the Functional Connectome after Chemogenetic Locus Coeruleus Activation.” Neuron.

